# Context matters: Situational stress impedes functional reorganization of intrinsic brain connectivity during problem solving

**DOI:** 10.1101/2020.05.26.117499

**Authors:** Mengting Liu, Robert A. Backer, Rachel C. Amey, Eric E. Splan, Adam Magerman, Chad E. Forbes

## Abstract

Extensive research has established the relationship between individual differences in brain activity in a resting state and individual differences in behavior. Conversely, when individuals are engaged in various tasks, certain task-evoked reorganization occurs in brain functional connectivity, which consequently can influence individuals’ performance as well. Here, we show that resting state and task-dependent state brain patterns interact as a function of contexts engendering stress. Findings revealed that when the resting state connectome was examined during performance, the relationship between connectome strength and performance only remained for participants under stress (who also performed worse than all other groups on the math task), suggesting stress preserved brain patterns indicative of underperformance whereas non-stressed individuals spontaneously transitioned out of brain patterns indicative of underperformance. These findings were subsequentially replicated in an independent sample set. Implications are discussed for network dynamics as a function of context.

---

Intrinsic patterns in functional connectivity predict a variety of meaningful diagnostic and behavioral outcomes (Krienen et al. 2014; Rosenberg et al. 2016; Tavor et al. 2016). At the same time, a host of literature documents that as people engage in cognitively demanding tasks, initial network components reorganize and become more specialized (Hugdahl et al. 2015; Schultz and Cole 2016; Bolt et al. 2017). Yet, to what extent network reorganizations occur from rest to task as a function of context, and the consequences this reorganization (or lack thereof) has for performance on cognitively demanding tasks remains unclear (Mill et al. 2017). The present study investigates how situationally stressful contexts alter the relationship between resting state and task-dependent brain network dynamics as they relate to performance on cognitively intensive tasks. Examining whole-brain connectivity patterns indicative of underperformance both before and during completion of a problem-solving task in either the presence or absence of stressors, we investigate whether context can dictate the link between specific brain patterns and underperformance by meaningfully altering transitions from rest to task; in this case, by inhibiting adaptive reconfiguration to task in a manner predictive of success on cognitively demanding tasks.

## Resting State and Task Dependent Brain Patterns are Associated with Optimal Cognition

Brain patterns at rest have been linked to a variety of task-related differences and cognitive performance capabilities, including emotion regulation (Morawetz C et al. 2016), attention (Rosenberg et al. 2016), distractibility (Poole et al. 2016), fluid intelligence (Finn et al. 2015), memory (Wang et al. 2010), and working memory (Magnuson et al. 2015). As such, resting-state networks are thought to reflect an underlying functional architecture that shapes network dynamics during cognitive processing (Wig et al. 2011; Mueller et al. 2013). As the brain undergoes transition from intrinsic to performance-related connectivity, intrinsic networks reorganize by recruiting different or additional regions to assist with specialized, functional processes (Cole et al. 2014). Moreover, the capacity to reorganize efficiently is important for successfully meeting the demands of a given task (Braun et al. 2015). Specifically, the ability to coordinate among *process-relevant* regions, while excluding noise from irrelevant regions, is ideal (Cole et al. 2013).

Problem solving is one such specialized functional process that is supported by a variety of functional neural components given its ultimate reliance on different executive processes, including updating or working memory, inhibition, and shifting (Miyake et al. 2000). Thus, it’s likely that both initial, intrinsic network properties and the ability to transition into alternate network configurations in response to task demands— configurations that instantiate successful execution of executive processes—collectively bear on successful outcomes (Anderson et al. 2014). For example, problem solving, intelligence, and visuospatial and working memory are all fortified by greater frontopareital network (FPN) cohesiveness, both at rest and during performance (Cocchi et al. 2013; Markett et al. 2014). In contrast, intrusion from other components during problem solving appears to interfere with FPN connectivity, undermining optimal processing. For instance, the activation of networks associated with self-referential and social cognition or emotion processing at times when people are attempting to problem solve predict worse performance, likely due to competition for limited cognitive and neural resources (Liston et al. 2009; Henckens et al. 2012; van Ast et al. 2016). This perspective is supported by recent findings indicating that stressed individuals’ suboptimal problem solving performance was characterized by less global neural network stability (Liu et al. 2017). This suggests that increased interactions between regions integral for emotion processing and regions integral for optimal performance on cognitively demanding tasks ultimately may be bad for performance as a whole.

## Context May Influence the Brain’s ability to Adaptively Transition between States

If intrinsic networks can influence task-related neural responses, then it stands to reason that they do so by either facilitating cognition that is congruent with demands of a task (when the task requires cognitive operations that are similar to what is already more intrinsically connected), or hindering cognition when the demands of a task conflict (e.g. if a performance context requires problem-solving, but also introduces evaluative stress, thus facilitating self, social, or emotion-oriented processes). In the case of the latter— hindered cognition—it’s possible that individual predispositions in resting connectivity result in maladaptive neural activation, which acts to reduce efficiency of adaptation to the ostensible task demands.

Borrowing from clinical literature, past research suggests that neural transitions from default mode network (DMN) dominant activity to FPN dominant activity is generally facilitated among depressed and anxious individuals, with the reverse observed among healthy individuals (i.e., in depressed subjects, FPN activation couples with more error-related rumination processing compared to DMN activation, and vice-versa for controls) (Hamilton et al. 2011). That is, the neural transition between brain states is dependent in part on both the context and individual differences in intrinsic brain patterns. The current study extends this work by examining whether a neural transition tendency from rest to task occurs for individuals differently according to their contexts, which in this case consists of a situation being more or less stressful.

Taken together, both intrinsic and task-dependent brain states are associated with performance, and stress may alter how intrinsic and task-related brain networks interact. Moreover, people vary in the extent to which they are susceptible to stressors, suggesting performance is affected by individual differences in resting network connectivity interacting with task-related context. Investigating intrinsic networks instantiating multiple psychological processes is inherently complex and thus traditional hypothesis-driven approaches may fail to identify unique patterns characterizing underperformance in stressful contexts. One promising approach to addressing this complexity is connectome predictive modeling (CPM) – a completely data driven approach that treats all possible pairs of brain regions and their associated connectivity values as individual, logistic predictors. CPM then constructs a model (the “connectome”) that maximally fits behavioral scores (Rosenberg et al. 2017; Shen et al. 2017). As such, CPM is a data-driven, predictive metric, which can be leveraged to identify what aspects of network properties across the whole brain characterize underperformance in individuals. Critically, connectivity provides a window into how different regions are functioning *in relation* to each other. As communication occurs rapidly across multiple timescales during problem-solving, we measured neural activity and connectivity using electroencephalography (EEG), which affords optimal temporal and frequency resolution that would be critical for disambiguating such complex relationships.

In this study, a connectome was first identified from whole-brain, resting-state connectivity that was predictive of underperformance across all individuals. This approach is analytically based on Rosenberg’s et al.’s approach but differs in that it was applied to a cross-frequency design, and targeted performance outcomes versus clinical diagnoses; hence, the term *connectome* is used to refer to the model. Connectivity values from those same regions identified at rest were then extracted from epochs of problem solving intervals where individuals solved math problems varying in difficulty in the presence or absence of situational stressors. Regarding network dynamics, the extent to which the connectome continued to predict performance when applied to *task*-related connectivity should reflect less network reconfiguration from rest to task, i.e., individuals fail to transition from a network state indicative of underperformance at rest to a network associated with optimal performance during the task. Regarding specific functional themes, past literature highlights the importance of attention, working memory, and executive control for successful math performance. However, according to van Ast et al. (2016), stressful performance contexts instead engender more emotion-related processing, at the expense of these functions. In particular, contexts that produce stress from priming evaluative threats also give rise to self and other-related cognition, as individuals remain vigilant for critical feedback about their behavior. Therefore, the data-driven network model should reflect related functional dynamics. Critically, however, if rest-to-task reconfiguration were compromised or inhibited due to situational stress, this pattern would linger for those under stress but not for their non-stressed counterparts. Moreover, the consequence of such a system disruption could possibly be observed in other, more broad network dynamics, such as global network efficiency. Thus, the connectome points to a possible key component in bringing about these more generally disruptive phenomena.

To address these premises, it is necessary that participants should not differ across confounding psychological dimensions (e.g. math anxiety) that would be associated with suboptimal cognitive functioning aside from context. One way to randomly select from a sample of cognitively normal individuals, ensure all individuals are exposed to the same stimuli, but place only one group uniquely in a stressful context is to prime stereotype-based stress (SBS) in women. SBS is a robust situational stressor that individuals experience as a result of primed context cues suggesting they may be negatively evaluated based on their membership in a stigmatized group, not unlike studies on social evaluative threat that utilize the common Trier Social Stress Test (Schmader et al. 2008; Hall et al. 2015; Forbes et al. 2018; Kirschbaum, Pirke, & Hellhammer, 1993; Allen et al. 2014). Consequently, this form of evaluative threat initiates a persistent cycle of hypervigilance and attempts to regulate accompanying emotions, both of which have been shown to interfere with cognitive capacity that is otherwise needed for optimal performance on cognitively intensive tasks (Schmader et al. 2008). As such, SBS provides a means to place one group under stress while holding other aspects of the situation and individual constant.

Utilizing two independent datasets (one to identify brain patterns associated with performance—i.e., Study 1—and one to replicate patterns identified in the initial dataset—i.e., Study 2) we collected resting state EEG data, then placed men and women in contexts that were either stress-free or uniquely stressful to only one group of women. Individuals then completed cognitively intensive, i.e., difficult math problems while continuous EEG activity was recorded. We hypothesized that to the extent stress may limit the likelihood that individuals’ transition from intrinsic network states at rest to network states more adaptive for cognitively demanding tasks, for women under stress (i.e., women in SBS contexts), intrinsic network markers of performance at rest would therefore continue to predict performance outcomes when applied to task activity, compared to non-stressed women and all men. This would possibly be due to interference with typical network reconfiguration otherwise normally evident during transitions from rest to problem solving. Conversely, to the extent the absence of stress allows for individuals to more easily transition from brain states optimal (or suboptimal) for performance at rest to performance during the task, individuals not under stress—and regardless of gender—should exhibit a network of regions predictive of performance *during* the task that differs from the network associated with performance among everyone at rest.

## Materials and Methods

### Participants

158 white participants (85 females) taken from a larger database (Forbes et al. 2018) completed this study for payment. We recruited only participants who were aware of the negative female-math stereotype. Specifically, participants needed to score a 3 or lower on the following question during a pre-study screening in order to qualify for the current study: “Regardless of what you think, what is the stereotype that people have about women and men’s math ability” (1= Men are better than women; 7= Women are better than men).” Seven participants were excluded in EEG analyses because their EEG data lacked one or more specific cognitive tasks (either resting or math solving tasks).

### Procedure

Upon entering the room, participants were taken to a soundproofed EEG chamber, seated in front of a computer and set up for EEG recording. Participants were randomly assigned to either the *stress condition* (i.e., eliciting stereotype threat-based stress for women) or the *control condition*. In the stress condition, participants were told that the results of the following tasks would be diagnostic of their math intelligence. In the control condition, participants were told that the results of the following tasks would be diagnostic of the types of problem-solving techniques they prefer (Forbes and Leitner 2014; Forbes et al. 2015). To further prime stereotype-based stress in the stress condition, participants marked their gender, sessions had a male experimenter present, and all instructions were read via a male experimenter’s voice (following protocol of Forbes et al., 2018). Conversely, participants in the control condition did not mark their gender, all sessions had all female experimenters present, and all instructions were read via a female experimenter’s voice. Following the instructions, participants completed a math feedback task for 34 minutes. Participants then answered a series of post-task questionnaires, were debriefed, and were paid for their participation.

### Resting state

Baseline EEG data was collected before the task began. Participants were asked to sit quietly in the EEG chamber, with directions to either sit resting with their eyes closed, or to blink normally. Baseline recordings were taken for five minutes, wich yields 175 eyes open EEG trials and 175 eyes close EEG trials Obtained EEG data (collapse both eyes open and eyes closed trials) from this time period constituted participant’s resting state brain activity.

### Math feedback task

Participants completed a 34-minute math task (Forbes et al., 2018). The task consisted of standard multiplication and division problems (e.g. 7×20=) that initial pilot tests confirmed varied in degree of difficulty (easy, medium and difficult, ensuring all participants would solve problems correctly and incorrectly). In a given trial, participants were provided with three answer options for each problem (A, B or C), with the answer to each problem randomly placed in one of the three answer positions. Participants then answered each multiple-choice problem using a button box placed on their laps, and were not permitted to use scratch paper. After entering their answer, participants received veridical feedback on the accuracy of their answer on their monitor for a duration of 2 seconds. After the feedback was presented, the next problem was presented on the screen. Each problem was presented for a maximum of 16 seconds. If participants didn’t complete a given problem within 16 seconds, they would receive negative feedback (i.e., that they got the problem wrong). On average, participants completed 83.9 problems. Dividing the total number of correct responses by the total number of attempted problems gave us our calculation for math score accuracy for both easy and difficult problems respectively.

### EEG measurement & preprocessing

#### Recording

Continuous EEG activity was recorded using an ActiveTwo head cap with an ActiveTwo Biosemi system (BioSemi, Amsterdam, Netherlands). Recordings were collected from 128 Ag-AgCl scalp electrodes and bilateral mastoids. Two electrodes were also placed next to each other one cm below the right eye to record startle eye-blink responses. A ground electrode was established using BioSemi’s common Mode Sense active electrode and Driven Right Leg passive electrode. EEG activity was digitized with ActiView software (BioSemi) and sampled at 2,048 Hz. Data was downsampled post-acquisition and analyzed at 512 Hz.

#### Preprocessing

For feedback analyses, the EEG signal was epoched and stimulus locked from 500ms pre-feedback presentation to 2,000ms post-feedback presentation. EEG artifacts were removed following FASTER (Fully Automated Statistical Thresholding for EEG artifact Rejection) (Nolan et al. 2010) preprocessing protocol—an automated approach to cleaning EEG data that is based on multiple iterations of independent component analysis (ICA) and statistical thresholding analyses. In FASTER, EEG trials are filtered by a 0.3-55 Hz band-pass FIR filter, and baseline-corrected using the time series from 100 ms preceding onset of tasks. First, EEG channels with significant unusual variance (operationalized as activity with an absolute z score larger than 3 standard deviations from the average), average correlation with all other channels, and Hurst exponent were removed and interpolated from neighboring electrodes using spherical spline interpolation function. Second, EEG signals were epoched and baseline corrected and epochs with significant unusual amplitude range, variance, and channel deviation were removed. Third, the remaining epochs were transformed through ICA. Independent components with significant unusual correlations with EMG channels, spatial kurtosis, slope in filter band, Hurst exponent, and median gradient were subtracted and the EEG signal was reconstructed using the remaining independent components. Finally, EEG channels within single epochs that displayed significant unusual variance, median gradient, amplitude range, and channel deviation were removed and interpolated from neighboring electrodes within those same epochs.

#### Source localization

Forward and inverse whole-brain models were calculated with an open access software, MNE-python (Gramfort et al. 2013; Gramfort A et al. 2014). The forward model solutions for all source locations located on the cortical sheet were computed using a three-layer boundary element model (BEM) (Hamalainen and Sarvas 1989) constrained by the default average template of anatomical MNI MRI. Cortical surfaces, extracted with FreeSurfer (Fischl 2012), were sub-sampled to about 10,240 equally spaced vertices on each hemisphere. The noise covariance matrix for each individual was estimated from the pre-stimulus EEG recordings after the preprocessing. The forward solution, noise covariance, and source covariance matrices were used to calculate the dynamic statistical parametric mapping (dSPM) estimated (Dale et al. 2000) inverse operator (Dale et al. 1999). Inverse computation was done using a loose orientation constraint (loose = 0.11, depth = 0.8) (Lin FH et al. 2006). The surface was divided into 68 anatomical regions of interest (ROIs; 34 in each hemisphere) based on the Desikan–Killiany atlas (Desikan et al. 2006). For each participant, a time course was calculated for each area/node by averaging the localized EEG signal of all of its constituent voxels at each time point during task performance.

### Functional connectivity estimation and network construction

Frequency coupling was calculated within identical frequency bands and temporal periods between all pairs of nodes. Phase locking values (PLV) (Lachaux et al. 1999), which measure variability of phase between two signals across trials, were utilized to define connectivity strength. In other words, for every participant, condition, and frequency band, we obtained a symmetric 68 × 68 adjacency matrix, representing all pairs of nodes—or *edges*—in each participant’s whole-brain network during a given period (Figure 1). For the math task period, PLVs were averaged from the first 1,000ms after the math problems appeared, because the first 1000ms involved a common time interval of all trials for each participant in solving math problems. For the resting state period, PLVs were averaged from the first 500ms after the onset of the initial fixation cross.

**Figure 1.**
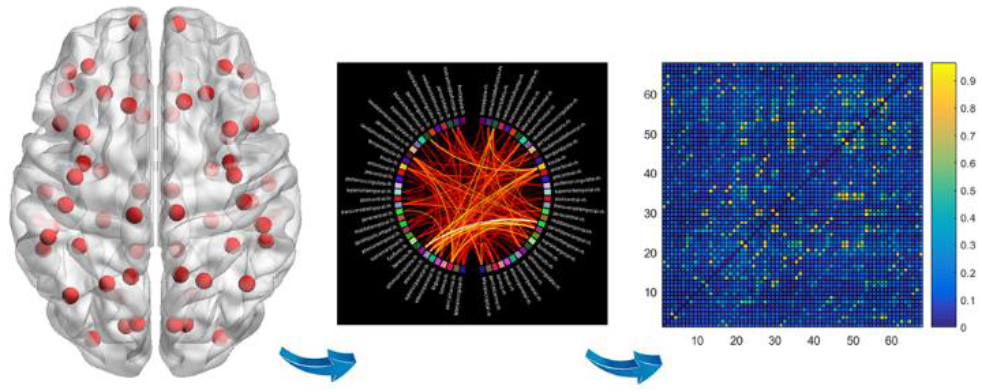
Construction of adjacency matrix. The cortical surface was divided into 68 anatomical regions of interest (ROIs) based on the Desikan–Killiany atlas (Desikan et al., 2006). Phase locking values were calculated to define the connectivity strength between all possible pairs of nodes. Finally, a symmetric 68 × 68 adjacency matrix was created to record all pairs’ connectivity strength.

### Prediction of functional connectivity to behavior

To assess the relevance of functional neural network connections to behavior, math scores (for both easy and difficult problems) were regressed on each of the resting-state network edges for all frequency bands, from *n*-1 participants. The resulting 2,244 x 4 = 8976 *p*-values for each regression were recorded in a 68 x 68 symmetrical matrix from 4 frequency bands, by participant. To identify significant edge associations and to minimize false positives stemming from multiple comparisons, a *p*-value threshold criterion of 0.001 was applied to all matrices. Thus, only pairs whose *p*-values were below 0.001 were retained as part of the network (Figure 2). The identified network was then utilized for the left-out participant in both rest and task state to test their predictive power to math performance. This procedure repeated *n* times via a leave-one-out cross validation.

**Figure 2.**
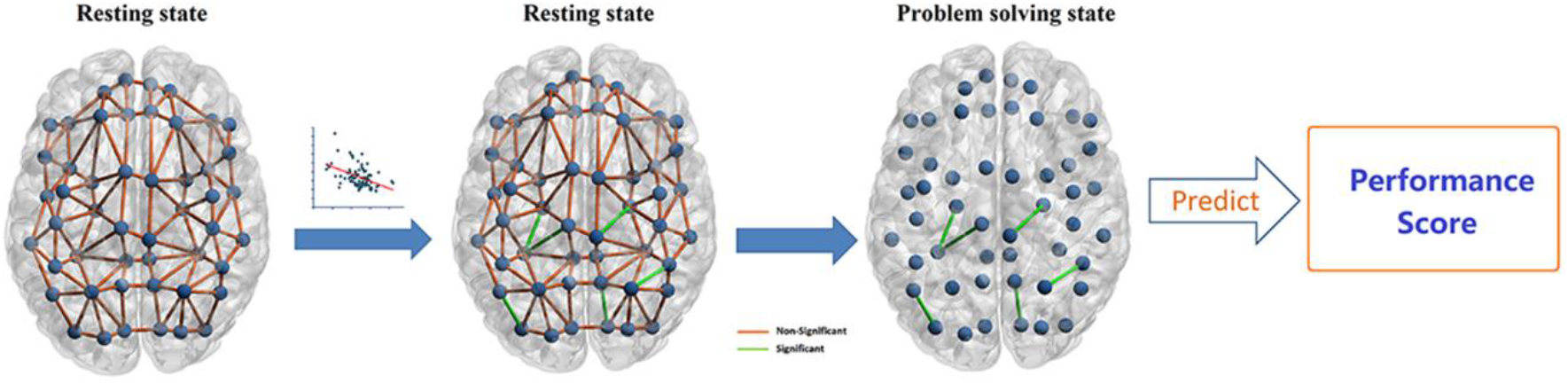
Simple linear regressions were conducted for each edge in the (resting state) connectivity matrices and math performance score, producing a matrix of pairs. The matrix was then thresholded based on *p* values, retaining a subset meeting significance criteria. Linear regression analysis was then likewise conducted for task-related connectivity and problem performance scores.

### Graph Theory Analysis of whole-brain functional networks

Two measures were chosen to explore the topography of the global functional matrices: *Clustering coefficient* and *global efficiency*. For group comparison, average network strength was also explored. Clustering coefficient measures represent the extent to which nodes in a graph tend to cluster together. This index provides a measure of local information transfer. Global efficiency is a measure of information transfer average across all the nodes. It’s been well established that the transition of brain networks from rest to task coincides with increases in clustering coefficients and global efficiency and the ability to accomplish complex cognitive tasks more robustly and efficiently (Bullmore and Sporns 2009; Bolt et al., 2017). Further information about these metrics can be found elsewhere (Rubinov and Sporns 2010). Graph theory analysis was performed using the Brain Connectivity Toolbox in MATLAB (https://sites.google.com/site/bctnet).

To sparse graphs, the whole-brain functional networks were thresholded using a range of density thresholds with any edge greater than the threshold set to “1” and any edge less than the threshold set to “0.” Density thresholds ranged from 5% (the minimum threshold where all graphs were fully connected) to 70% in 5% increments. Using multiple definitions of the graph model ensured that group differences were not dependent on arbitrary factors such as thresholds (Scheinos et al., 2017; Garrison et al. 2015).

## Results

### Manipulation check: Stress and Math performance score

Initial analyses on amygdala activity (assessed via startle probes elicited to positive and negative feedback and operationalized as a measure of stress) were conducted in Forbes et al. (2018). These analyses indicated that all participants elicited marginally greater amygdala responses to negative feedback received on the math feedback task compared to positive feedback and women elicited larger amygdala responses to feedback compared to men. However, only women in the stress condition exhibited a unique non-linear (quadratic) relationship in their amygdala responses to feedback over time, suggesting a unique stress response among individuals experiencing evaluative threat. Women in the stress condition also performed worse on the math task compared to all other conditions, a typical finding in the evaluative threat literature (planned contrast on math test accuracy: t (1, 156)=3.17, p=.002, d=.51). These behavioral results overall provide supporting evidence that the evaluative threat and thus stress induction manipulation was successful (see details in supplementary results).

### Network prediction of performance

To clarify relationships between edges and performance (connections that had positive or negative relationships with performance), edges were further grouped into two separate networks by the valence of their *R*-values. By averaging all edge’s *PLV values* within each of the networks, a summary statistic—*network strength*—characterized each participant’s degree of positive and negative connectivity. Math scores were then regressed on to the “negative” and “positive” network strength values, for both easy and difficult problems (all *p*s < 0.001).

To determine whether these networks—originally defined from resting period data—remained predictive when applied to task-related activity, math scores on easy and difficult problems were, again, each regressed on network strength of the same two positive and negative networks (i.e., based on the same edges at rest) *during the task period* using leave-one-out cross validation.

The negative network remained predictive of performance for difficult problems during task activity (*β* = 0.25, *F*(1,149)=8.32, *p* = 0.004). All other relationships were non-significant during task activity (all *p*s > 0.22; Figure 3A). That is, brain regions collectively exhibiting an increase in activation were associated with decreased performance on difficult (i.e., the most cognitively intensive) problems specifically.

**Figure 3.**
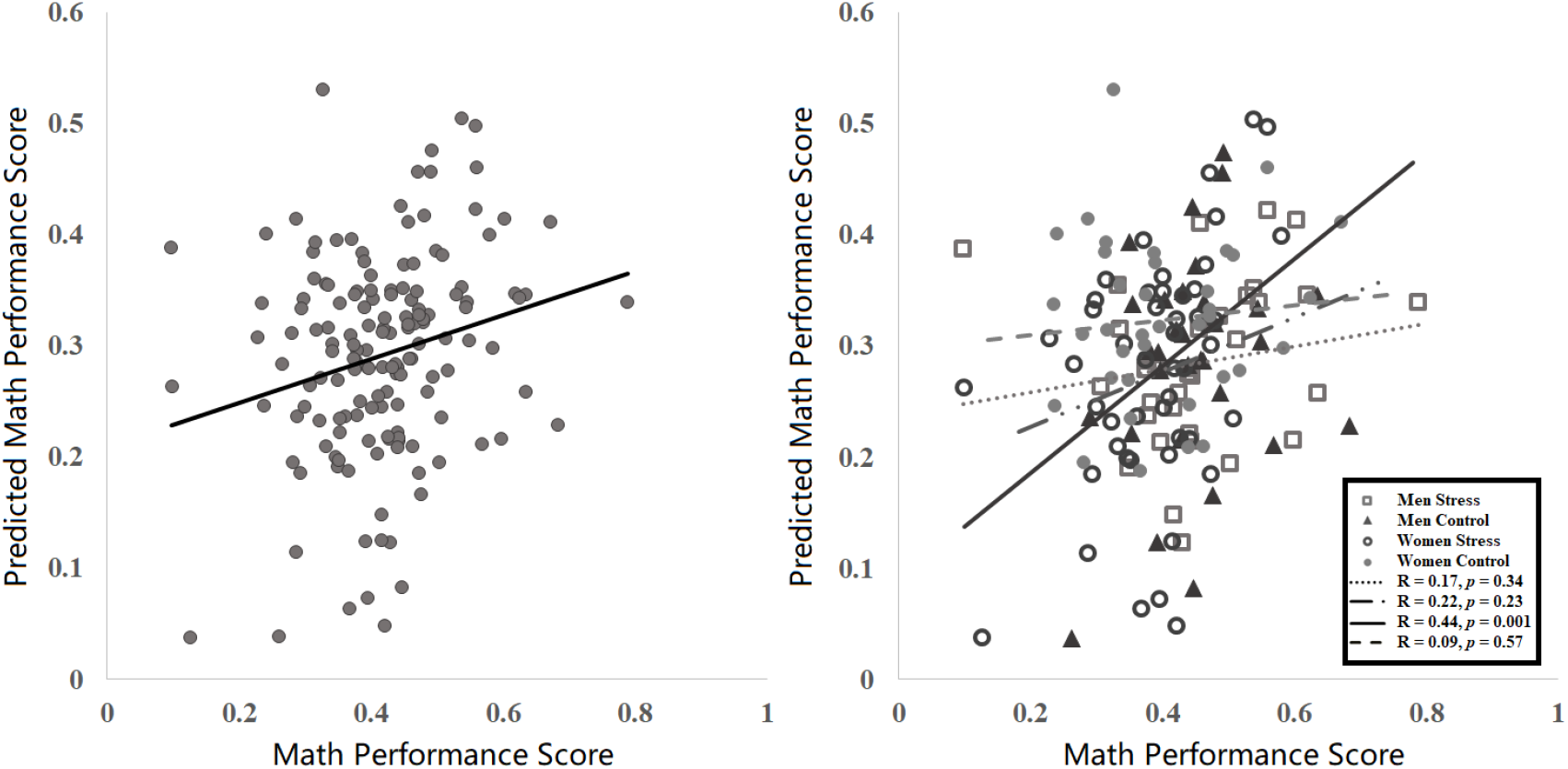
A) The same network identified in resting-state activity, when applied to task-related activity, was also predictive of math performance scores using cross-validation. This effect was significant only for performance on difficult math problems. B) The negative relationship between network connectivity and performance was driven by women under situational stress. This effect was not found in any other experimental conditions.

### Contextual influence

Next, we sought to determine whether context, and thus stress, played a role in the predictive relationship between task-related network strength and performance. Double moderation analyses assessed network strength’s predictive relationship with math scores (in the previously identified network from resting state), as a function of condition and gender (i.e., the moderators). These analyses used unstandardized regression coefficients and 95% bias-corrected confidence intervals (CIs) from 10,000 bootstrap estimates (Hayes 2013; model 3). 95% CIs are considered significant if the interval (e.g., [.3,.7]) does not contain zero (Cumming 2008). Results revealed a significant interaction between gender and condition (*p* =.0384, 95%, CI = [0.015, 0.52]). Conditional effects identified that, only for women in the stress condition, increased activation of the negative network during solving difficult problems predicted decreased performance (*β* = 0.54, *F*(1, 47) = 10.48, *p* =.0004,). Conversely, no such relationships were found among PST (i.e. control condition) women, as well as men in both conditions (all ps >.23), suggesting that the relationship between the negative network during task and performance outcomes, which was initially derived via resting state analyses, only persisted when women were experiencing stress (Figure 3B).

### Relevance of functional connectivity to behavior in non-stressed groups

If non-stressed individuals can spontaneously transition from intrinsic network properties to more adaptive network properties associated with optimal performance, then the next question is what are these network states? To answer this, we conducted an additional CPM analysis on task-related EEG activity for everyone but women in the stress condition. These analyses yielded positive and negative networks that significantly predicted participant’s math performance during the math task (*p*’s < 0.001).

To determine whether these network-behavior relationships—originally defined from task period data—were either evident intrinsically (i.e., remained predictive when applied to resting state activity) or re-organized during the rest to task transition (i.e., were not predictive in resting state activity), math scores of difficult problems were regressed on to network strength of these edges (i.e., based on the same edges at task) during the *rest* period. Results revealed both positive and negative networks were not predictive of performance for difficult problems during resting state activity (*p*’s > 0.382; Figure 4). This suggests that edges identified as predictive of performance during the task were the product of functional re-organization from rest to task for non-stressed participants.

**Figure 4.**
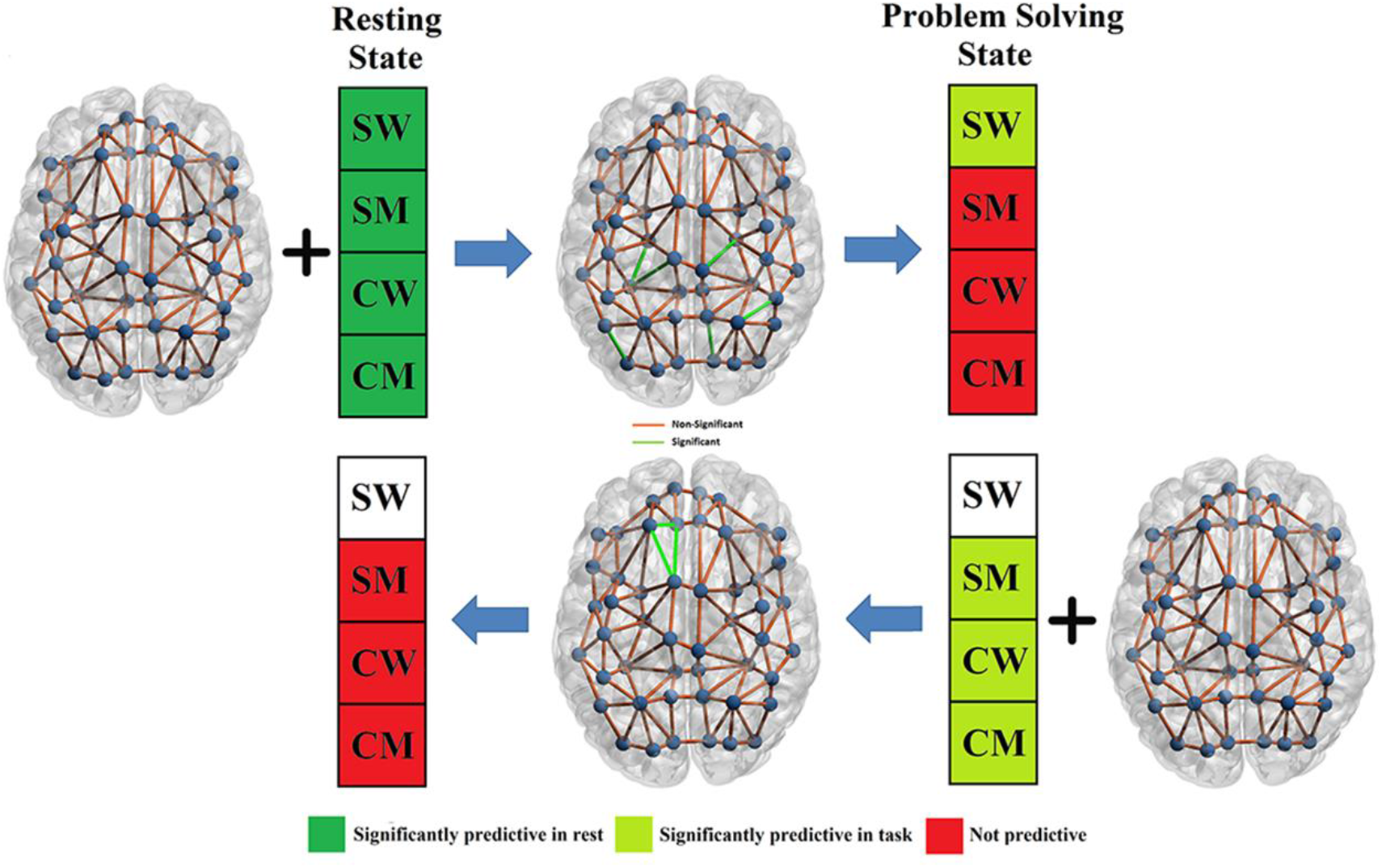
Predicting underperformance across experimental cells. At rest, the “bad” pattern is predictive of underperformance to varying degrees across all individuals. Still at rest, subdividing individuals by experimental cell, we observe the same pattern. During task, the bad pattern remains predictive (i.e., relevant to performance) only for individuals in the cell comprising women under threat. For non-stressed participants, i.e, control condition participants and men in the stress condition, functional connectivity patterns predictive of success were identified during problem solving epochs, which were not found to be predictive of performance in rest epochs. This suggests that non-stressed participants exhibited a functional state transition associated with optimal problem solving.

### Time variant relevance of functional connectivity to behavior

We found that the identified connectivity to math underperformance remains only for women in stress condition. However so far, we only demonstrate this in the first one second during the math solving process. Next, we explored whether this uninterrupted connectivity-behavior relation was more pronounced during a specific problem solving period, or was a more pronounced phenomena across the entirety of a problem solving epoch. To do this, a 5,000ms epoch was extracted from each problem-solving interval, and the 5,000ms epoch was spliced into 1,000ms time windows with an overlap of 500ms. Given that there is only minor difference between three non-stressed groups (see Figure 3), we collapse the participants from 3 non-stressed groups. Linear regression was conducted on observed math performance and predicted math performance and was compared between women in stress condition and non-stressed participants. Significant effects were found between 0-2500ms (p’s < 0.043, after multiple comparisons corrections) for women under stress (Figure 6A-B). No significant effects were found (p’s > 0.29) for other groups across the epoch. These findings indicated that the maladaptive state (with the functional connectivity predict under-performance) only emerge in the beginning of the problem-solving stage. The time course of R values across groups is plotted in Figure 6C.

**Figure 5.**
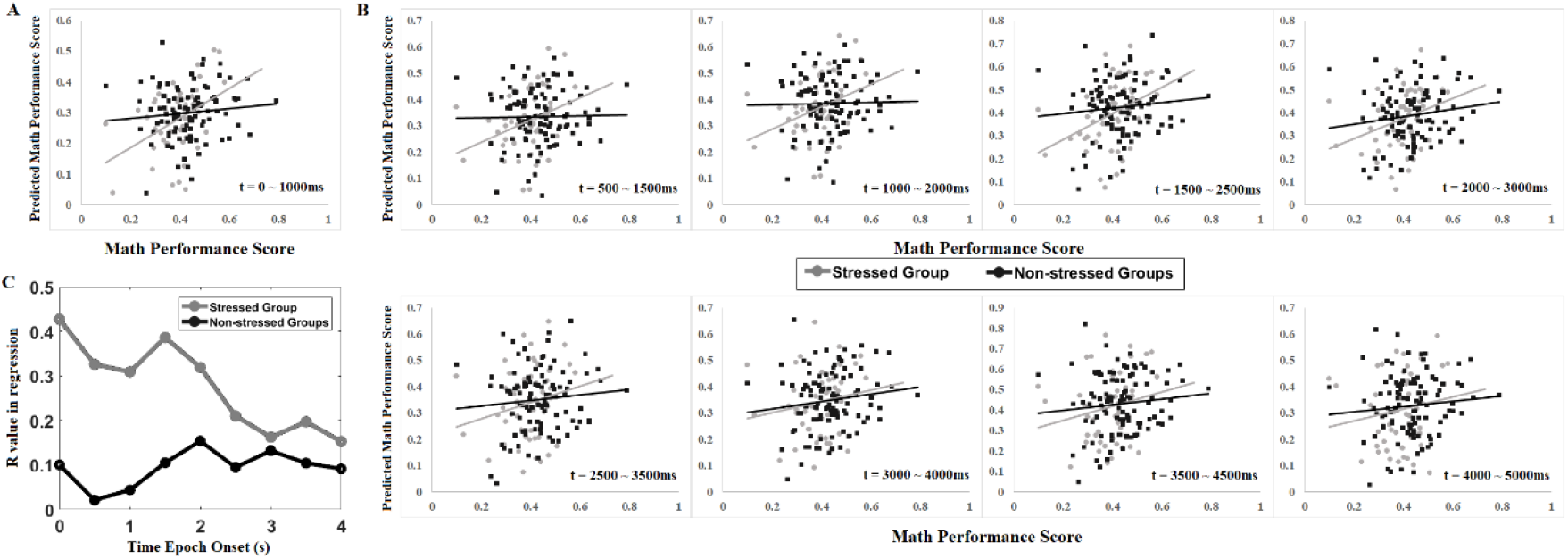
CPM analysis was extended up to 5000ms during math problem solving using a sliding window strategy, with a window length of 1 second and a moving step of 500ms. A) Math performance prediction using cross-validation based on functional network found in resting state (gray: women under stress; black: non-stressed participants). B) Math performance prediction within 1000ms window sliding from 500ms to 5000ms. C) Track of R values of regression curves for all of the sliding windows, x axis represent the start time for the sliding window. Significant effects for women under stress were found for first 4 sliding windows (0-2500ms).

**Figure 6.**
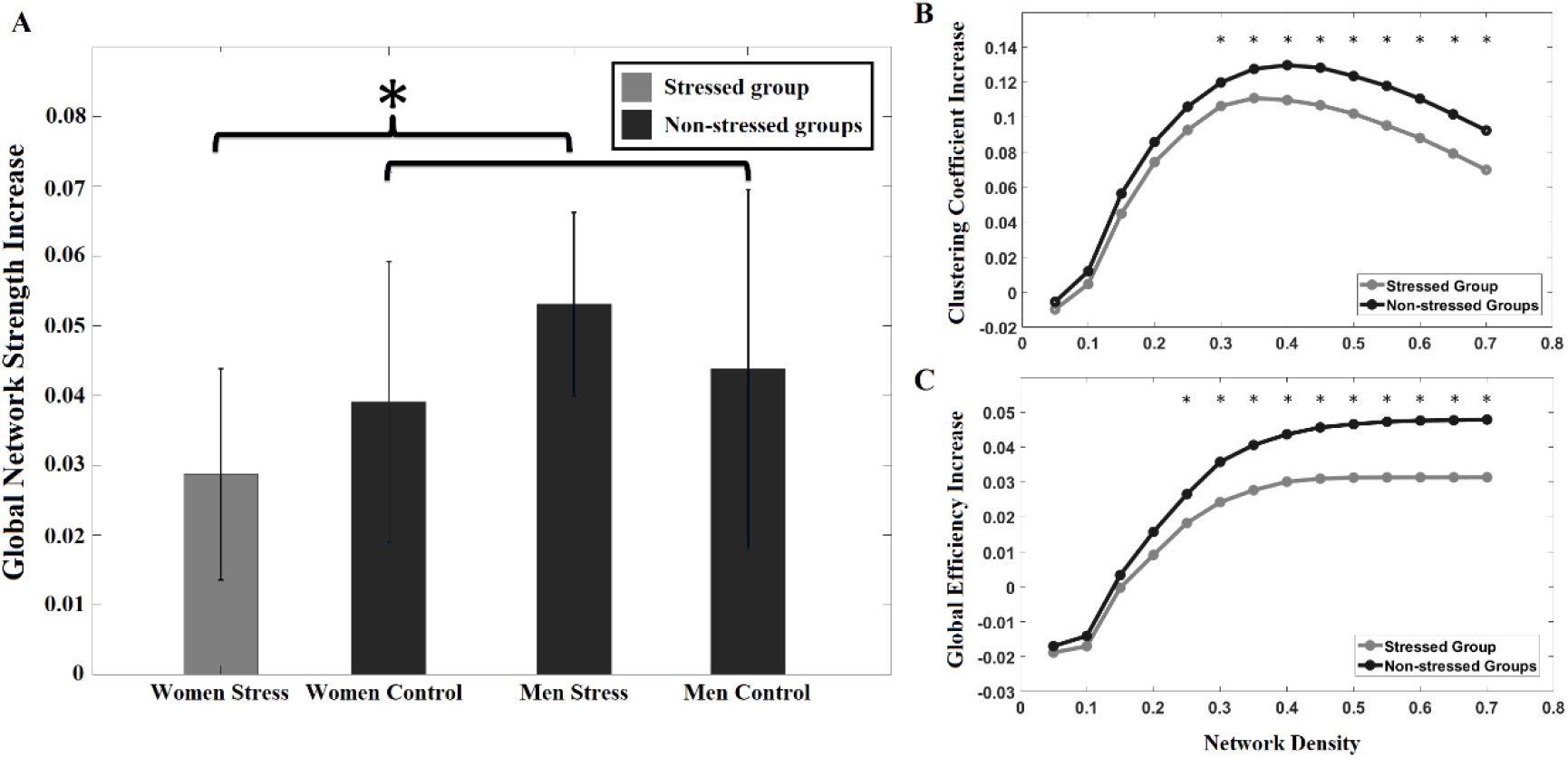
Functional network reorganization comparison between stressed group (women under stress) and non-stressed group (other three groups) in theta band. A) Women under stress showed less global network strength increase compared to non-stressed groups. B) women under stress showed a less increased clustering coefficient, as well as a less increased global efficiency compared with other groups.

### Direct comparison of functional connectivity organization alterations between stressed vs. non-stressed groups

We have provided evidence that the predictive power of identified connectivity to math underperformance only remains for women in stress condition. This implies that stress may facilitate the connectivity stability or attenuate the functional reorganization for optimal math performance. However so far, we only demonstrate the connectivity stability in regard to behaviors. To further verify that stress may have impeded functional network reorganization, we next examined different metrics of functional network organization regardless of behaviors. Specifically, we investigate the alteration of average connectivity, global efficiency and clustering coefficient (the two graph theory variables) for whole brain functional network, from resting state to the problem solving state (during the problem solving task) for women in the stress condition and other non-stressed groups. Given that the networks identified via CPM were specific to connectivity within theta and beta frequency bands, we yoked analyses involving global efficiency and clustering coefficient to these two frequency bands as well. Figure 5 represent the comparison of global connectivity alteration, and global graph theory variable alteration between women under stress and other non-stressed groups.Two way t-test was conducted on global average connectivity, global efficiency and clustering coefficient between women in stress condition and participants collapsed from all other three non-stressed groups. Results revealed that in the theta band, women in the stress condition exhibited significantly less increase of average PLV between regions (i.e., “strength”) compared with other groups (*p* = 0.022). In addition, after correcting for multiple comparisons, women under stress showed a less increased clustering coefficient averaged across all nodes in the graph, as well as a less increased global efficiency compared with other groups. Similar relationships were not evident in the beta band (Figure S2). These findings suggest that, regardless of math performance, functional connectivity alteration and functional network re-organization were attenuated by stress.

### Replication of current findings

To further validate the reliability of our findings, identical functional connectivity models were applied to an independent data set consisting of EEG data collected two years after Study 1, where participants completed a shortened version of the math feedback task described in Study 1. Results reaffirmed the presence of the negative network, which predicted math performance across participants in resting state, but remained predictive during the task only for women in the stress condition (see supplementary materials for details).

### Utilizing Neurosynth to explore potential functions of identified brain regions

Having determined that intrinsic neural patterns interacted with context to predict performance, we considered what functional themes might characterize, in particular, worse performance (i.e., the negative network). Assessing the specific functional roles of brain regions identified via data-driven approaches is a frequent challenge in neuroscience, one that is often susceptible to reverse inference biases. To attenuate this as best as possible, we conducted an exploratory meta-analysis using the *Neurosynth* program to outline potential functional roles of the regions identified from CPM analyses (http://neurosynth.org; Yarkoni et al. 2011). Given coordinates of a given brain voxel, Neurosynth provides terms most often associated with the area, ranked by posterior probability (*pp*) and *z* scores.

Examining the negative network at rest, three edges’ activation during the task (identified at rest) were found to predict poorer performance for all participants in all the cross-validation trials. The first edge consisted of connectivity between *right supramarginal gyrus* (R-SMG) and *right cuneus* (R-CUN) in theta band. The second edge consisted of connectivity between *right pars-opercularis* (inferior frontal gyrus; R-IFG) and *left pre-central cortex* (L-PreC) in beta band. The third edge consisted of connectivity between *right supramarginal gyrus* (R-SMG) and *left superior-parietal cortex* (L-SPC) in beta band. Note that based on moderated regression analyses, these findings were ultimately driven by women in the stress condition.

Evaluative threats have been described to undermine performance by initiating a cycle of hypervigilance and negative arousal, which is theorized to detract from requisite executive function resources (Schmader et al. 2008). We found some evidence of correspondence with these general themes in Neurosynth results; nodes of the negative network were associated with terms related to social cognition, emotion, and attention. For instance, R-IFG was strongly associated with “social” and “empathy” across both *pp* and *z* scores, as was L-SPC for “spatial attention”. Other frequent terms included “TOM” (theory of mind), “intentions”, and “mental state”, as well as “visual” and “spatial attention”. We observed the broad trend that edges appeared to reflect links between functions that revolve around internally oriented processing (self or social cognition; Amey et al., 2018) and those that typically relate to problem-solving (visuospatial attention, calculation, and responding; e.g. Anderson et al. 2014). However, these findings are exploratory in nature and as such, relationships remain tentative. Some discrepancies existed between *pp* and *z* scores (e.g. “emotion” and “pain” presented in *z* scores but not *pp*). We reserve further consideration about edge relationships, as well as full details regarding both networks, for the supplementary section.

## Discussion

Using a completely data-driven approach, findings from these studies a) identify intrinsic neural connectivity patterns that modulate problem-solving performance, but linger only in contexts facilitating evaluative stress, b) identify neural connectivity markers that predict performance among non-stressed individuals during a task but not at rest, and c) completely replicate across two large, independent datasets. Specifically, in Study 1, initially across *all* participants at rest, connectivity in three network edges were found to have a negative relationship with performance on difficult (i.e., more cognitively demanding) math problems: R-SMG and L-SPC, R-IFG and L-PreC, and R-SMG and L-SPC. During task-related activity, these connections remained predictive only for those in a stressful context (i.e. women primed with gender-salient, evaluative threat cues) solving difficult problems. Conversely, during the task, all other participants exhibited different neural connectivity patterns predictive of performance (Arsalidou et al. 2018), suggesting these individuals transitioned in to different brain states during the task. This suggests that appropriate task-reconfiguration necessary for optimal performance may be hindered by context, where in this case the experience of stress preserves maladaptive connectivity to modulate performance accordingly.

To further appraise the transition from rest to task, we evaluated the functional network structures regardless of their relationship to behavior. Stress context participants showed significantly reduced global connectivity increases between brain regions compared to other groups of participants. A global increase or decrease in functional connectivity from rest to task has not been observed consistently in past studies (Lynch et al. 2018). It may depend on tasks and even frequency bands (Ghaderi et al. 2019). Although functional connectivity during task cannot be regarded as the simple summation of functional connectivity at rest and task-evoked activity (Mastrovito 2013; He et al. 2013; Lynch et al. 2018), from a graph theoretical perspective, enhanced global connectivity may suggest an overall increased interconnection among the brain’s “hubs” (Scheinost et al. 2017). More integration of these hubs could thus support efficient communication between specialized regions across the brain in the service of better performance. Less enhanced global connectivity under conditions of stress may therefore reflect poorer network reorganization to meet task demands.

Additional evidence of a disruption in functional reorganization was also obtained from direct graph theory analyses, where stress context participants displayed less increased global efficiency and clustering coefficients. Generally, neural networks are expected to reconfigure from a modular, baseline topology at rest to a more efficient but costly topology during task (Meunier et al. 2010; Bertoleroa et al. 2015). That is, temporarily stronger direct connections between specialized nodes required for the task facilitate more efficient signal transmission. Thus, networks with high cluster coefficient and high global efficiency are advantageous to information processing during task (i.e. they reflect a “small-world” architecture; Basset and Bullmore 2006). Taken together, the stress group’s display of lower global efficiency and clustering supports that network transition was indeed impeded, irrespective of the functional properties of the network’s individual edges.

Another interesting consideration is whether the observed deficit of functional re-organization for stressed participants was stable across the entire problem solving process or just a transient state. Our time variant analysis (Fig. 6) revealed that the maladaptive patterns only predominated in the beginning of the problem solving process. That is to say, stressed participants may “shake off” this counterproductive state as time goes on. However, considering that the women under stress solved math problems less accurately than other groups, it indicates that stress nevertheless exerts influence on problem-solving abilities several seconds later. For instance, evaluative stress may evoke additional, processes that would be counterproductive to problem solving, e.g. emotion regulation or attempts to counter perceived critical feedback. Future studies are needed to further assess the nature of stressed participants’ states during later time points.

Examining further the “bad” pattern regions identified, three—L-PreC, R-CUN, and L-SPC—play integral roles within networks that facilitate successful math performance: the fronto-pareital (FP) executive and visuospatial attention (VSA) networks (Klingberg et al. 2002; Chambers and Prescott 2010). On the other hand, R-SMG and R-IFG are implicated in internally-oriented processes, like self and other cognitions or emotion regulation, which often detract from performance.

One interpretation is that the pattern could reflect a predisposition *towards* integration of these processes that, when cued by the context, would be inherently antagonistic for optimal math performance (van Ast et al. 2016). Alternatively, the pattern could represent a particular configuration within the whole-brain network at rest, which individuals formally transition *away* from, to math-solving related states, but failed to do successfully under stress (Fig. 7). These two explanations do not necessarily conflict. Future studies ideally would test which hypothesis most realistically reflects the mechanism underlying current findings.

Regardless of mechanism, the present findings are nonetheless intriguing. Past research demonstrated that intrinsic network states (Mennes et al. 2011; Zou et al. 2013; Rosenberg et al. 2016) and functional task reconfiguration (Schultz and Cole 2016) play integral roles in cognition and behavior. The present studies suggest that the role of these factors is dependent, to an extent, on the context in which these phenomena transpire. In other words, context may function to regulate certain neural adaptations, or selectively facilitate/impede *aspects* of adaptation. As such, the “bad” pattern could be regarded as a unique form diathesis that would otherwise not manifest when not elicited by these unique situational characteristics. The interpretation that stress hindered transition to more appropriate task-configurations is further supported by examining what everyone other than women in the stress condition transitioned to—presumably, connectivity facilitating better math performance. Indeed, in all experimental cells except the stressed group, greater fronto-parietal and visual spatial (externally oriented) connectivity and less internally-oriented connectivity predicted better performance (see supplemental materials), which is consistent with a number of past studies (Gerlach et al. 2014).

In order to hold all aspects of the context and individual constant while placing one group under evaluative stress, we employed a stereotype-based stress (SBS) paradigm to evoke evaluative threat in female subjects. Stress is a subjectively negative experience, accompanied by a cascade of chemical reactions that can adversely affect performance. In evaluative stress paradigms (e.g. the Trier stress test), stress elicits a host of maladaptive symptoms: increases in cortisol, alpha amylase, increased blood pressure and skin conductance, increased vACC and amygdala activity in response to stressors (negative feedback), and accompanying negative perceptions (Allen et al., 2014). SBS is unique in that only a derogated group experiences the evaluative stress (Schmader, Johns, & Forbes, 2008). Thus, SBS and more domain-general evaluative threats like TST are synonymous in symptoms, differing only in the pervasiveness of the stressor. Some evidence finds that men may experience higher cortisol in stress paradigms, though women report higher negative psychological experiences (Liu et al., 2017). Other work has also argued women may be more prone to specifically math-related anxiety (Maloney, Waechter, Risko, & Fugelsang, 2012). However, given that only the women who underwent the SBS manipulation displayed elevated stress and worse performance, we find it reasonable to expect that findings from the present paradigm would generalize to other populations for whom an evaluative threat is salient.

Several findings were difficult to interpret definitively in the current study and remain open questions for future studies. For instance, it is unclear why the link between intrinsic connectivity patterns and underperformance in stressful contexts were specific to difficult math problems and not easy math problems. One possible explanation for these findings stems from past evaluative threat oriented research. Past work on evaluative threat/stereotype based stress clearly demonstrates that threat-based underperformance is typically most pronounced on the most cognitively intensive tasks. In fact, when individuals experiencing SBS complete easier problems or specific types of tasks where success is dependent on prepotent responses, their performance is often facilitated (Spencer et al. 1999; Jamieson et al. 2007). This suggests that contexts need to be more stressful to preserve intrinsic network properties’ influence during performance, or keep individuals “stuck” in a maladaptive network state; however, future research would be necessary to validate this theory.

Future work will enhance our understanding of the present phenomena in several ways. The CPM approach employed in this study (by virtue of our research question) identified key network markers, derived from phase-locking averaged across time and trials during two temporally generalized events: rest and performance. Further research should clarify how these features realistically interact with other key regions and normal brain states, as parts of process-relevant networks. This would be particularly helpful for understanding the full picture of the rest-task brain network reorganization.

Moreover, this novel CPM approach could be applied to additional contexts, where a similar relational perspective stands to provide insight. Aside from stress, other contexts likely elicit unique reorganization from intrinsic networks to task activation—for example, situations requiring multitasking may favor intrinsic markers that either downplay conflicts, or aid in supporting simultaneous/parallel processes, or the integration of various processes. Hence, the ability to compare predictive markers across various states and contexts could provide a means to obtain greater dimensionality in functional themes.

While the EEG-based approach utilized in these studies provided high quality temporal resolution to capture these phenomena, a measure of caution is warranted when interpreting activation as regionally specific, due to the inherent spatial limitations of the methodology. By using a high-density electrode array, adopting advanced Bayesian source localization (incorporating inverse dSPM operators), and engendering a more conservative estimation of neural activity by constraining to regions in proximity of the brain surface using MNE, it is possible to make better inferences about the roles of specific brain regions (Cohen 2014). However, due to lack of precise individual-level brain forward modeling, EEG source localization may only be accurate to the precision of a few centimeters (Cohen 2014). Hence the functional interpretation of the mechanism behind our findings remain tentative. Future research should complement these findings by applying spatially attuned, fMRI imaging—particularly given the importance of amygdala and medial temporal regions in arousal and self-referential processes (Forbes et al. 2018). Another limitation is the number of participants in each group is relatively small (n < 50), especially in replication study. This may have a reduced chance of detecting a true effect and can increase the false positive rate (Button et al. 2013; Yarkoni, 2009). Future effort will be made by recruiting more participants and a power analysis will be conducted before a study begins.

## Conclusion

Employing a CPM approach to characterize markers of cognitive performance and manipulating evaluative stress across contexts, findings across two independent datasets revealed that contexts can modulate optimal or suboptimal neural reconfigurations from rest to task. Stress rendered predisposed individuals “stuck” in maladaptive patterns, whereas non-stressed individuals tended to transition into brain states associated with optimal performance, illustrating one way in which task-evoked adaptation is contingent on contextual demands. This predisposition was reflected on cognitively intensive problems in particular. By illustrating how unique network characteristics provide a means for individuals to thrive (or falter) in specific contexts, these findings draw closer to more ecologically valid models of cognition—and, hopefully, more fruitful, brain-based stress interventions moving forward.

## Acknowledgements

All aspects of this study and article were supported by National Science Foundation grants #1329281 and #1535414 awarded to Chad E. Forbes.

---

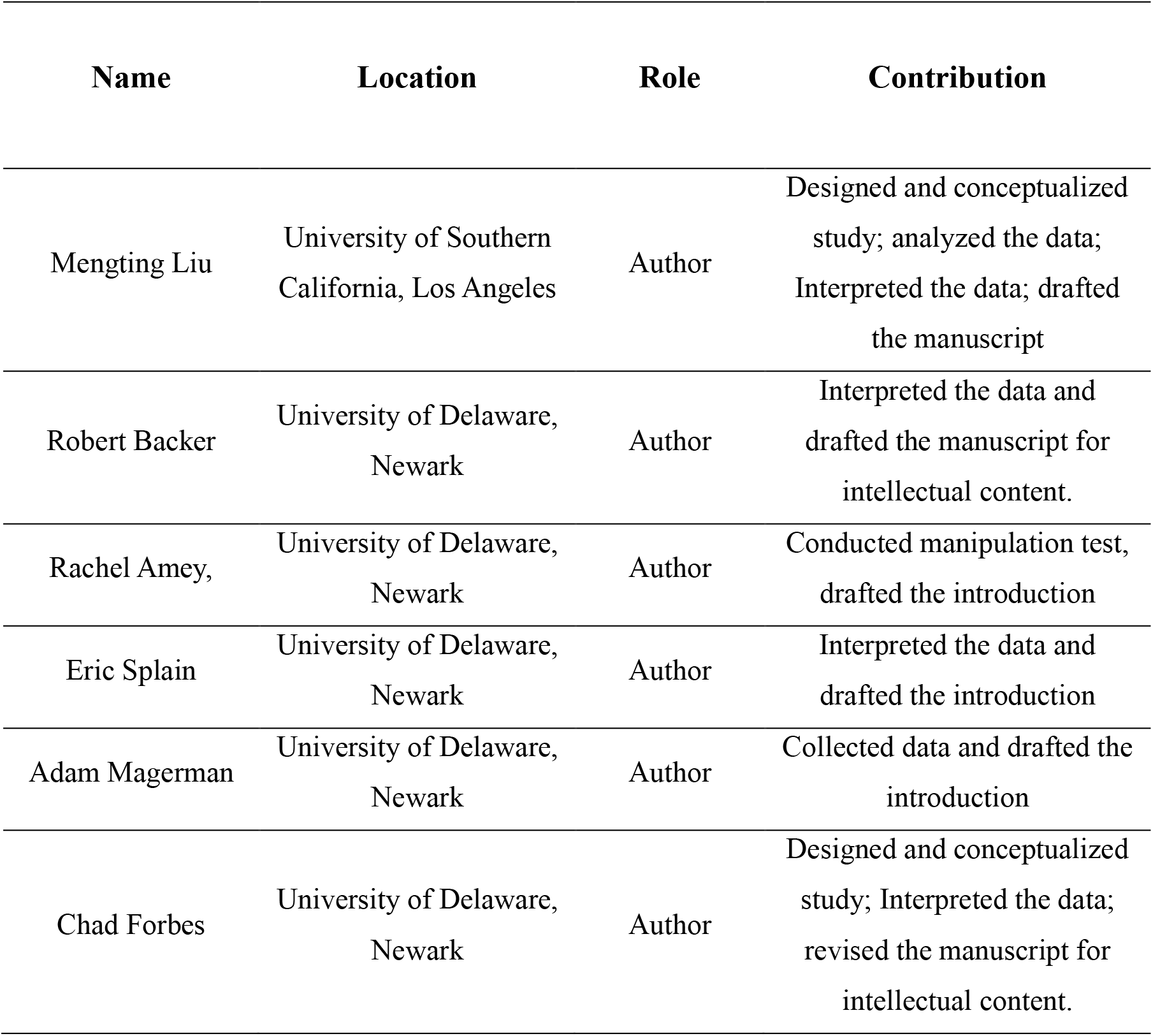
Authors’ contributions

## Supplemental Information

### Stress manipulation check: Stress and math performance score

To examine whether there were overall performance differences between conditions on math score performance, an initial 2 (Gender: Men or Women) x 2 (Condition: DMT or PST) factorial ANOVA was conducted on participants’ accuracy on the math feedback task (number correct/number attempted). This analysis yielded a main effect for gender, *F* (1, 156) =16.56, *p*<.001, *d*=.63. There were no other main effects or interaction (*p*’s>.38). Given the well documented effects of stereotype threat on performance (for a review see Schmader et al., 2008), however, planned contrasts were also conducted to compare DMT women’s performance on the math feedback task to the other three conditions. These analyses indicated that DMT women (i.e., those experiencing stereotype threat) performed worse on the math feedback task compared to the other three conditions, *t* (1, 156)=3.17, *p*=.002, *d*=.51.

With respect to the role of task difficulty, given the nature of our math feedback task, which was lengthy and contained easy, medium and difficult questions, it’s possible that these various problem types had variable effects on performance across groups. To examine this, a 2 (Gender: Men or Women) x 2 (Condition: DMT or PST) x 3 (Problem Type: easy, medium or difficult) mixed factors ANOVA with repeated measures on the latter variable was conducted. This analysis yielded a main effect for gender, *F*(1, 156)=12.38, *p*=.001, *d*=.55, that was qualified by a problem type by condition interaction, *F*(1, 156)=4.44, *p*=.01, *d*=.35, and problem type by gender interaction, *F*(1, 156)=3.41, *p*=.03, *d*=.29. Simple effects analyses using a Dunn-Sidak adjustment to control for multiple comparisons indicated that DMT women performed worse on easy, F(1, 156)=4.17, p=.04, d=.33, and difficult problems, F(1, 156)=4.73, p=.03, d=.35, compared to PST women. Men did not differ from one another with respect to condition, p’s>.23. DMT women also performed worse on easy, F(1, 156)=11.99, p=.001, d=.56, and moderately difficult problems, F(1, 156)=3.77, p=.05, d=.31, compared to DMT men. Women in the PST condition performed worse on easy, F(1, 156)=6.51, p=.01, d=.41, and moderate problems, F(1, 156)=5.64, p=.02, d=.38, compared to men in the PST condition. Interestingly only DMT women did not perform differently on easy, medium and difficult problem types, presumably because they underperformed across problem types in general (p’s>.12). All other groups showed the expected patterns, i.e., performing better on easy compared to moderate and difficult problems, and moderate compared to difficult problems (p’s<.04). Results overall provide supporting evidence that the stereotype threat manipulation was successful.

### Study 2—a replication of our findings

In study 2 to an independent data set consisting of EEG data collected from 101 white participants (60 females) two years after Study 1 who completed a shortened version of the math feedback task described in Study 1. As in Study 1, participants were only recruited if they were aware of the negative female-math stereotype. 11 participants were excluded because EEG collected was significantly contaminated with artifacts during at least one of the cognitive tasks. Instructions and priming procedures were identical to Study 1, with the exception that participants completed a shortened, 17-minute version of the original math feedback task (see Supplemental Methods for further information). Thus, this was a 2 (Condition: stress or control) by 2 (Gender: Men or Women) factorial between-subjects design. Participants also completed a series of tasks that were used as part of a separate study. EEG measurement, preprocessing, source localization and network construction were identical to Study 1.

### Performance on the math feedback task for replication study

An initial 2 (Gender: Men or Women) x 2 (Condition: DMT or PST) factorial ANOVA was conducted on participants’ accuracy on the math feedback task (number correct/number attempted). This analysis yielded a main effect for gender, *F* (1, 100) =27.04, *p*<.001. There were no other main effects or interaction (*p*’s>.41). Given the well documented effects of stereotype threat on performance (for a review see Schmader et al., 2008), however, planned contrasts were also conducted to compare stereotype threatened women’s performance on the math feedback task to the other three conditions. These analyses indicated that women under stereotype threat performed worse on the math feedback task compared to the other three conditions, *t* (1, 100)=-3.32, *p*=.001, *d*=.64.

### Brain-behavior relation for replication study

Using a completely data-driven CPM approach, findings from Study 1 identified connectivity properties within a number of regions and specific frequency that predicted performance. For women in stressful contexts, the same connectivity patterns that predicted performance at rest predicted performance during the task, specifically on the most cognitively intensive math problems (the most difficult problems). These same patterns did not predict performance for individuals in stress-neutral contexts. Instead, a different collection of brain regions, and connectivity between these brain regions specifically, predicted performance during the task. Given the nature of our data-driven approach, the question remains as to what extent these findings are ultimately meaningful, i.e., are these findings replicable? To test the reliability of our data-driven findings, we applied the network models identified in Study 1 to an independent data set consisting of EEG data collected two years after Study 1 who completed a shortened version of the math feedback task described in Study 1.

Identical analyses to Study 1 were conducted: EEG activity elicited during problem solving was isolated and connectivity between *R-IFG* and *L-PreC* and *R-SMC* and *L-SPC* in beta band, as well as *R-SMG* and *R-CUN* in theta band, was calculated. The summation of these connections was then used to predict performance on difficult problems at rest for all participants, as well as on the math task (i.e., the most cognitively intensive problems where ST effects are typically most pronounced) as a function of gender and condition in a double moderated regression analysis. Consistent with findings from Study 1, in resting state, results revealed a significant relationship between summation of negative network strength and performance on the difficult math problems (*β* = −0.48, *p* = 0.013). Examining network strength between these regions during the task, results yielded a marginally significant relationship between the summation of negative network edges and performance on difficult math problems (*β* = −0.37, *p* = 0.031); all other relationships (for easy problems and positive network in difficult problems) were non-significant (*p*’s > 0.32; Figure S1A). Furthermore, an interaction between gender and condition was also replicated (*p* =.043); as with Study 1, only women in the stress condition performed worse on difficult math problems to the extent that the negative network (identified originally at rest) was better connected during the solving of difficult math problems (*B* = -.3033, *p* =.0043, 95% CI [-.6718, -.0651]). Women in the control condition and men in both conditions again showed no relationship among these variables (*ps* >.61; Figure S1B), confirming once more that the relationship between the network (identified both at rest and during performance) and performance on difficult math problems was evident only when individuals were experiencing stress.

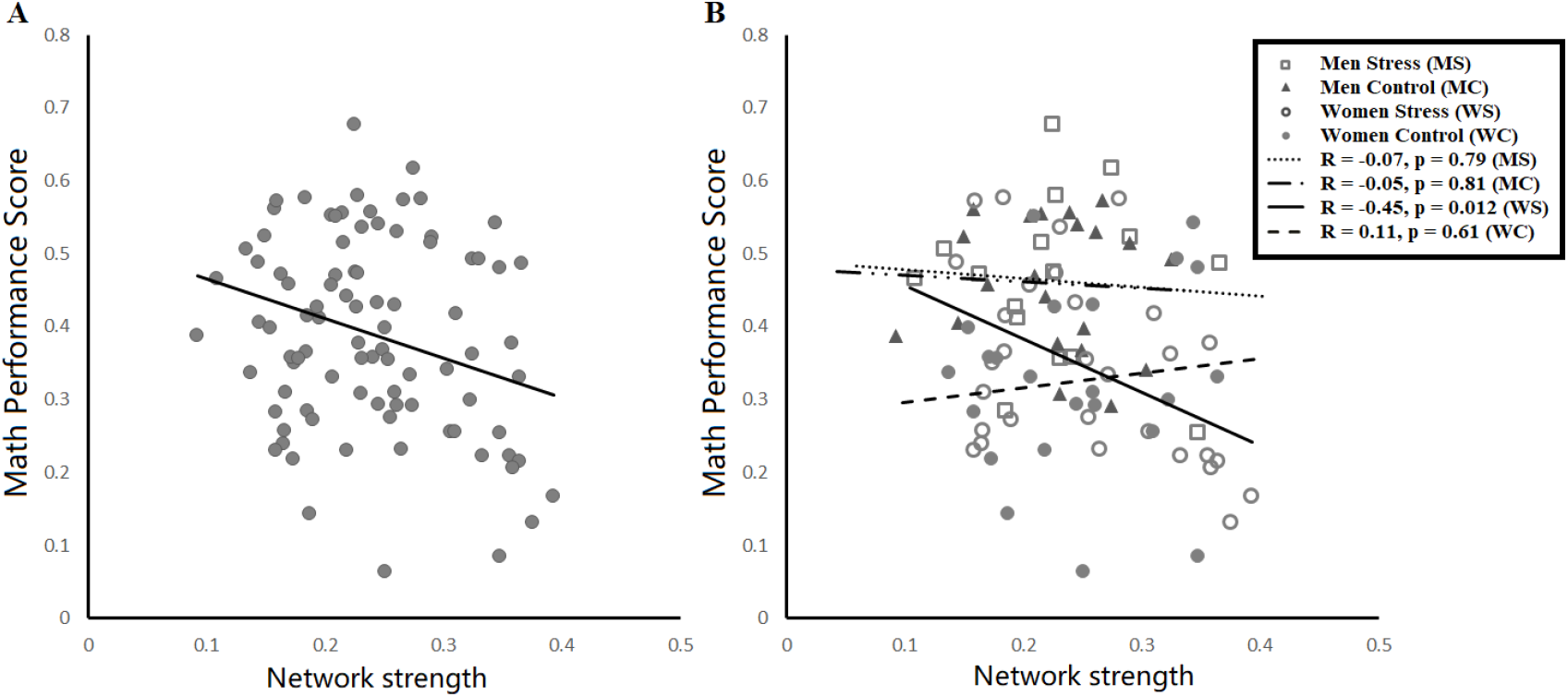

### Direct comparison of functional connectivity organization alterations between stressed vs. non-stressed groups in beta band

**Figure S2:**
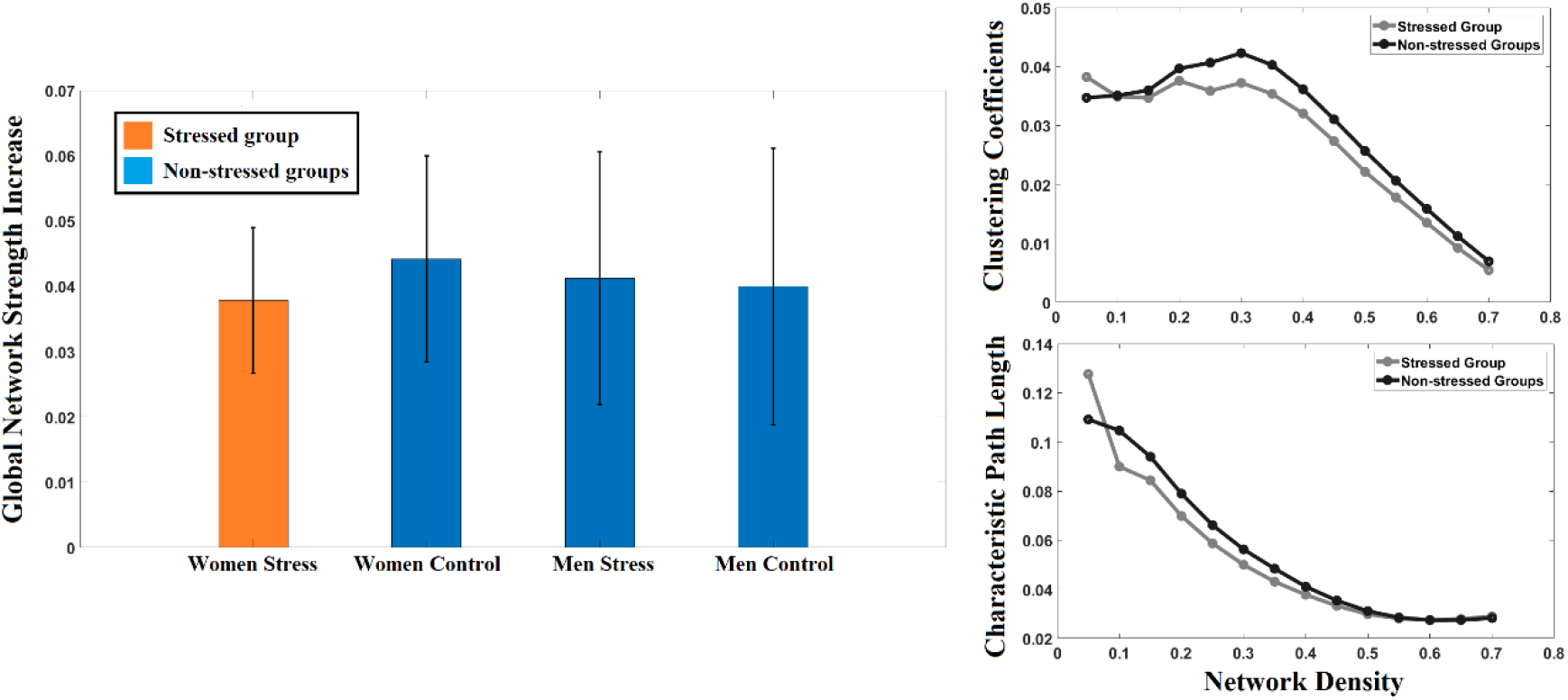
Functional network comparison between stressed group (women under stress) and non-stressed group (other groups) in beta frequency band. No significant differences were found.

### No contextual bias in resting state

Before making the conclusion that people went through a brain state transition process from rest to problem solving except for DMT women, we need to further validate that the neural-performance found in resting state for all participants is not driven by DMT women. That is the relationship for other three groups were also successfully established, and they truly processed a brain state transition. To examine this, linear regression was conducted on network strength and math performance in non DMT-women participants in resting state. Results revealed significant effects for all other three groups (p < 0.001), suggesting no contextual bias for the relationship between the negative network in resting state and math performance. These results suggested that only women in the DMT condition (the only stressed group) maintains their brain-behavior relationship during the transition from resting state to task state. Other participant experience a brain functional reorganization.

### Relaxing the Threshold for Data-Driven Networks

To check whether the results were merely a product of our initial, stringent *p*-value threshold, we increased the threshold to 0.005 and 0.01 respectively. Connectivity values whose *p*-values were below the two thresholds in resting state were retained. In math solving tasks, the PLV values within a given network were summed and averaged. Linear regression analyses were conducted on the identified positive and negative networks for easy and hard math problems, respectively. Results indicated that the significant effect was only kept (*p* < 0.001, *r* = 0.289 for threshold of 0.005; *p* < 0.001, *r* = 0.301 for threshold of 0.01) when solving hard math problems for negative network.

Double moderated regression was also conducted on dataset with increased threshold. Condition and gender were utilized as moderators. We tested for double moderation by deriving unstandardized regression coefficients and 95% bias-corrected confidence intervals (CIs) from 10,000 bootstrap estimates (Hayes, 2013; model 3). For threshold of 0.05, results revealed a marginal significant interaction between gender and condition (*p* =.0727). Conditional effects identified that, only for women in the stress condition, increased activation of the negative network during solving difficult problems predicted decreased performance (*β* = 0.29, p =.002, 95%, CI = [-0.021, 0.18]). Conversely, no such relationships were found among PST (i.e. control condition) women, as well as men in both conditions (all ps >.09). For threshold of 0.01, analyses revealed that no significant effect for gender, condition or gender x condition, indicating that effects of selected functional connectivity on performance are not moderated by stereotype-based stress. However, conditional effects revealed that for DMT women only, performance score for hard problems during problem solving correlated to predicted performance score for hard task (B=0.213, p=.009). Other groups showed no relationship (p’s>.15). Results suggested that participants in different group have same effects, but women under stress has the strongest.

### Functional Decoding of Identified Brain Regions

Results (Fig. S3) reveal that R-IFG and R-supramarginal cortex were primarily associated with emotional processing, with some relation to social applications. For example, they are closely linked to terms such as empathy, pain, distractor, emotion, inhibition and social. R-IFG was also found to be involved in speech related terms such as tones, sentence, and R-supramarginal cortex was found involved in motor and imagine, which are less clearly related to our math solving task. L-precentral cortex was strongly linked to motor related terms such as motor, premotor and supplemotor. L-precentral was also associated with terms of eye and execution, which relates to visuospatial attention. In addition, the term somatosensory could be reflect part of the precentral area that is close to postcentral cortex, which is thought to be the primary somatosensory cortex. L-SPC is mostly related to spatial sense related terms, such as spatial attention, direction and action observation. However, L-SPC is was also found to be related to execution related terms such as working memory (WM), fronto-parietal network, and execution. These associated terms are reasonable because such functions are thought to be involved in posterior-parietal cortex, which could be an extension of SPC. Overall, given the experimental context of solving math problems in stressful situation, functional connectivity between R-IFG and L-precentral could be interpreted as more association between emotional regulatory and visuospatial processing. Function connectivity between R-supramarginal cortex and L-SPC, on the other hand, can be viewed as more association between social inference and visual orienting processes (which we may go so far as to call “vigilance”).

**Figure S3:**
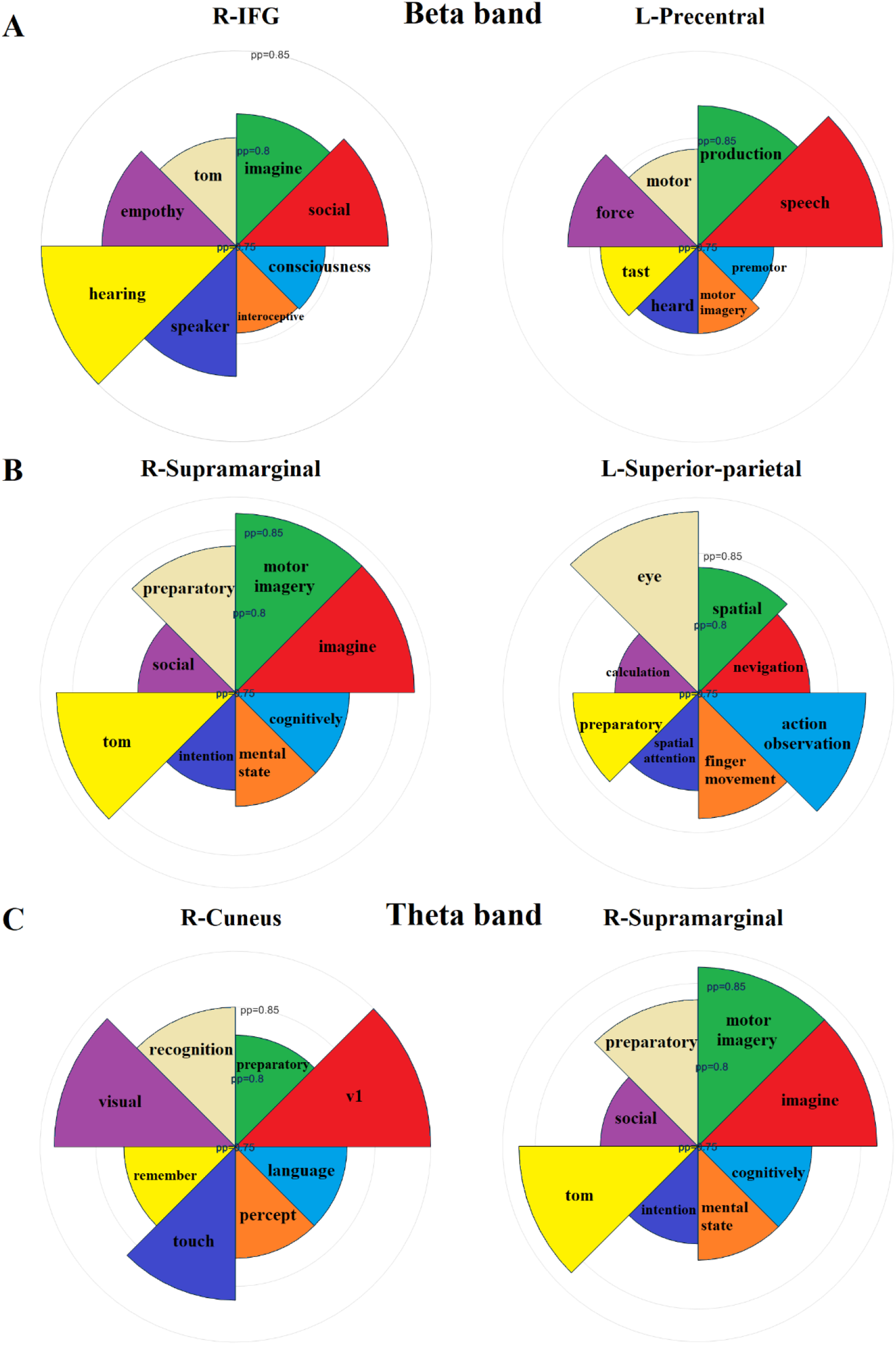
Functional decoding of regions between negative edges in resting state for all participants. Functional terms were identified by the data-driven model, provided by Neurosynth. Eight of the most highly associated cognitive functions are selected, according to their inverse reference posterior probability (pp), for each pair of region: A) *right pars-opercularis* (inferior frontal cortex) and *left pre-central cortex* connectivity, B) *right supramarginal cortex* and *left superior-parietal cortex* and e) *right cuneus* and *right supramarginal cortex*.

